# Integrating healthcare and research genetic data empowers the discovery of 28 novel developmental disorders

**DOI:** 10.1101/797787

**Authors:** Joanna Kaplanis, Kaitlin E. Samocha, Laurens Wiel, Zhancheng Zhang, Kevin J. Arvai, Ruth Y. Eberhardt, Giuseppe Gallone, Stefan H. Lelieveld, Hilary C. Martin, Jeremy F. McRae, Patrick J. Short, Rebecca I. Torene, Elke de Boer, Petr Danecek, Eugene J. Gardner, Ni Huang, Jenny Lord, Iñigo Martincorena, Rolph Pfundt, Margot R. F. Reijnders, Alison Yeung, Helger G. Yntema, DDD Study, Lisenka E. L. M. Vissers, Jane Juusola, Caroline F. Wright, Han G. Brunner, Helen V. Firth, David R. FitzPatrick, Jeffrey C. Barrett, Matthew E. Hurles, Christian Gilissen, Kyle Retterer

## Abstract

*De novo* mutations (DNMs) in protein-coding genes are a well-established cause of developmental disorders (DD). However, known DD-associated genes only account for a minority of the observed excess of such DNMs. To identify novel DD-associated genes, we integrated healthcare and research exome sequences on 31,058 DD parent-offspring trios, and developed a simulation-based statistical test to identify gene-specific enrichments of DNMs. We identified 285 significantly DD-associated genes, including 28 not previously robustly associated with DDs. Despite detecting more DD-associated genes than in any previous study, much of the excess of DNMs of protein-coding genes remains unaccounted for. Modelling suggests that over 1,000 novel DD-associated genes await discovery, many of which are likely to be less penetrant than the currently known genes. Research access to clinical diagnostic datasets will be critical for completing the map of dominant DDs.

## Introduction

It has previously been estimated that ~42-48% of patients with a severe developmental disorder (DD) have a pathogenic *de novo* mutation (DNM) in a protein coding gene^1,2^. However, over half of these patients remain undiagnosed despite the identification of hundreds of dominant and X-linked DD-associated genes. This implies that there are more DD relevant genes left to find. Existing methods to detect gene-specific enrichments of damaging DNMs typically ignore much prior information about which variants and genes are more likely to be disease-associated. However, missense variants and protein-truncating variants (PTVs) vary in their impact on protein function^3–6^. Known dominant DD-associated genes are strongly enriched in the minority of genes that exhibit patterns of strong selective constraint on heterozygous PTVs in the general population^7^. To identify the remaining DD genes, we need to increase our power to detect gene-specific enrichments for damaging DNMs by both increasing sample sizes and improving our statistical methods. In previous studies of pathogenic Copy Number Variation (CNV), utilising healthcare-generated data has been key to achieve much larger sample sizes than would be possible in a research setting alone^8,9^.

## Improved statistical enrichment test identifies 285 significant DD-associated genes

Following clear consent practices and only using aggregate, de-identified data, we pooled DNMs in patients with severe developmental disorders from three centres: GeneDx (a US-based diagnostic testing company), the Deciphering Developmental Disorders study, and Radboud University Medical Center. We performed stringent quality control on variants and samples to obtain 45,221 coding and splicing DNMs in 31,058 individuals (**Supplementary Fig. 1**; **Supplementary Table 1**), which includes data on over 24,000 trios not previously published. These DNMs included 40,992 single nucleotide variants (SNVs) and 4,229 indels. The three cohorts have similar clinical characteristics, male/female ratios, enrichments of DNMs by mutational class, and prevalences of known disorders (**Supplementary Fig. 2**).

To detect gene-specific enrichments of damaging DNMs, we developed a method named DeNovoWEST (*De Novo* Weighted Enrichment Simulation Test, https://github.com/queenjobo/DeNovoWEST). DeNovoWEST scores all classes of sequence variants on a unified severity scale based on the empirically-estimated positive predictive value of being pathogenic (**Supplementary Fig. 3-4**). We perform two tests per gene: the first is an enrichment test on all nonsynonymous DNMs and the second is a test designed to detect genes likely acting via an altered-function mechanism. This second test combines an enrichment test on missense DNMs with a test of linear clustering of missense DNMs within the gene. We then applied a Bonferroni multiple testing correction accounting for 18,762 × 2 tests, which takes into account the number of genes and two tests per gene.

We first applied DeNovoWEST to all individuals in our cohort and identified 281 significant genes, 18 more than when using our previous method^1^ (**Supplementary Fig. 5**; **Fig. 1a**). The majority (196/281; 70%) of these significant genes already had sufficient evidence of DD-association to be considered of diagnostic utility (as of late 2019) by all three centres, and we refer to them as “consensus” genes. 54/281 of these significant genes were previously considered diagnostic by one or two centres (“discordant” genes). Applying DeNovoWEST to synonymous DNMs, as a negative control analysis, identified no significantly enriched genes (**Supplementary Fig. 6**).

**Figure 1:**
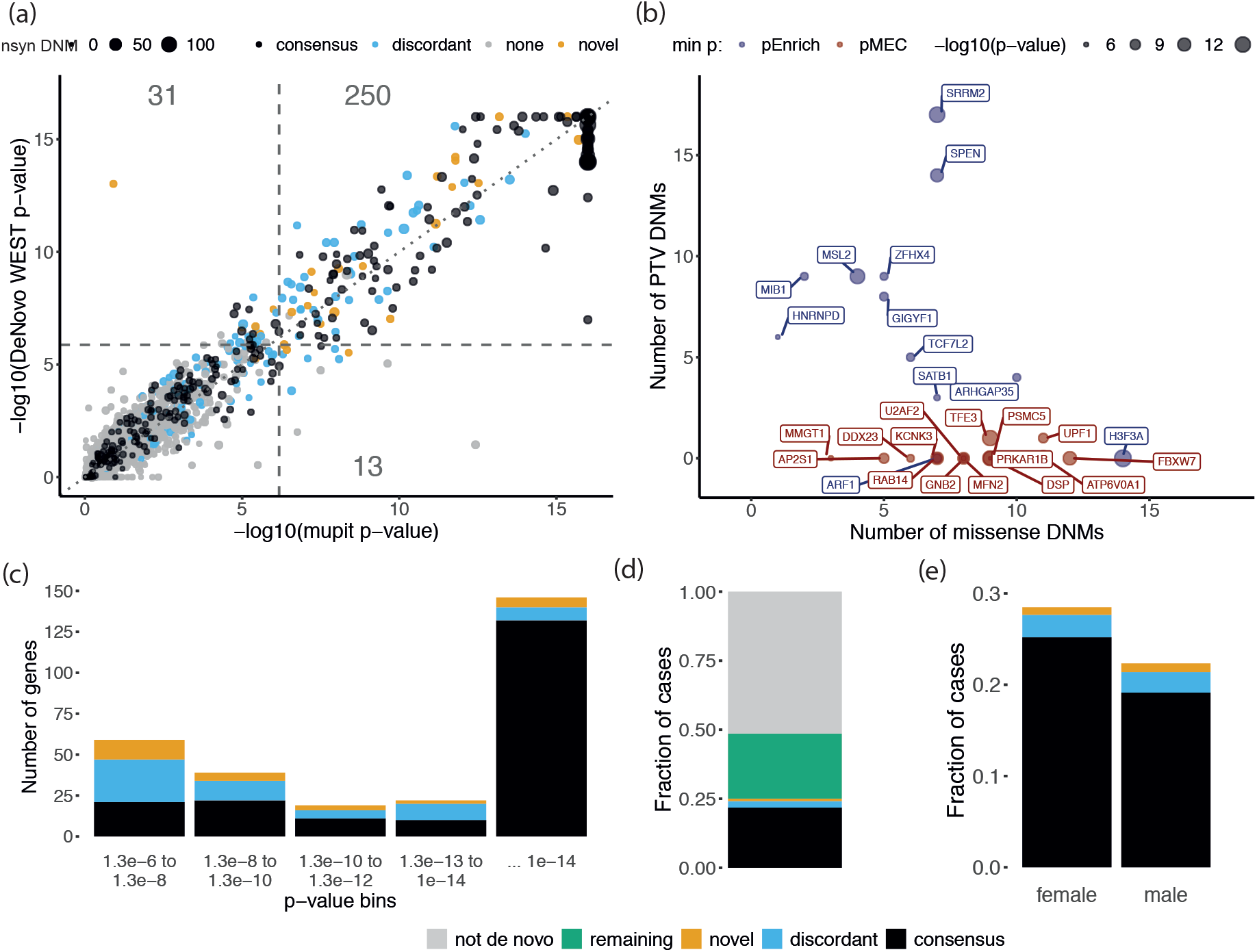
Results of DeNovoWEST analysis. (a) Comparison of p-values generated using the new method (DeNovoWEST) versus the previous method (mupit)^1^. These are results from DeNovoWEST run on the full cohort. The dashed lines indicate the threshold for genome-wide significance. The size of the points is proportional to the number of nonsynonymous DNMs in our cohort (nsyn). The numbers describe the number of genes that fall into each quadrant (b) The number of missense and PTV DNMs in our cohort in the novel genes. The size of the points are proportional to the log_10_(-p-value) from the analysis on the undiagnosed subset. The colour corresponds to which test p-value was the minimum (more significant) for these genes: non-synonymous enrichment test in blue (pEnrich), or missense enrichment and clustering test in red (pMEC). (c) The distribution of p-values from the analysis on the undiagnosed subset for discordant and novel genes; p-values for consensus genes come from the full analysis. The number of genes in each p-value bin is coloured by diagnostic gene group. (d) The fraction of cases with a nonsynonymous mutation in each diagnostic gene group. (e) The fraction of cases with a nonsynonymous mutation in each diagnostic gene group split by sex. In all figures, black represents the consensus genes, blue represents the discordant genes, and orange represents the novel genes. In (c), green represents the remaining fraction of cases expected to have a pathogenic *de novo* coding mutation (“remaining”) and grey is the fraction of cases that are likely to be explained by other genetic or nongenetic factors (“not *de novo*”).

To discover novel DD-associated genes with greater power, we then applied DeNovoWEST only to DNMs in patients without damaging DNMs in consensus genes (we refer to this subset as ‘undiagnosed’ patients) and identified 94 significant genes (**Fig. 1b; Supplementary Fig. 7; Supplementary Table 2**). While 61 of these genes were discordant genes, we identified 33 putative ‘novel’ DD-associated genes. To further ensure robustness to potential mutation rate variation between genes, we determined whether any of the putative novel DD-associated genes had significantly more synonymous variants in the Genome Aggregation Database^5^ (gnomAD) of population variation than expected under our null mutation model (Supplementary Note). We identified 11/33 genes with a significant excess of synonymous variants. For these 11 genes we then repeated the DeNovoWEST test, increasing the null mutation rate by the ratio of observed to expected synonymous variants in gnomAD. Five of these genes then fell below our exome-wide significance threshold and were removed, leaving 28 novel genes, with a median of 10 nonsynonymous DNMs in our dataset (**Fig. 1c**; **Supplementary Table 3**). There were 314 patients with nonsynonymous DNMs in these 28 genes (1.0% of our cohort); all DNMs in these genes were inspected in IGV^10^ and, of 198 for which experimental validation was attempted, all were confirmed as DNMs in the proband. The DNMs in these novel genes were distributed approximately randomly across the three datasets (no genes with p < 0.001, heterogeneity test). Six of the 28 novel DD-associated genes are further corroborated by OMIM entries or publications, including *TFE3*^11,12^ for which patients were described in two recent publications.

We also investigated whether some synonymous DNMs might be pathogenic by disrupting splicing. We annotated all synonymous DNMs with a splicing pathogenicity score, SpliceAI^20^, and identified a significant enrichment of synonymous DNMs with high SpliceAI scores (≥ 0.8, 1.56-fold enriched, p = 0.0037, Poisson test; **Supplementary Table 4**). This enrichment corresponds to an excess of ~15 splice-disrupting synonymous mutations in our cohort, of which six are accounted for by a single recurrent synonymous mutation in *KAT6B* known to disrupt splicing^21^.

Taken together, 25.0% of individuals in our combined cohort have a nonsynonymous DNM in one of the consensus or significant DD-associated genes (**Fig. 1d**). We noted significant sex differences in the autosomal burden of nonsynonymous DNMs (**Supplementary Fig. 8**). The rate of nonsynonymous DNMs in consensus autosomal genes was significantly higher in females than males (OR = 1.16, p = 4.4 × 10^−7^, Fisher’s exact test; **Fig. 1e**), as noted previously^1^. However, the exome-wide burden of autosomal nonsynonymous DNMs in all genes was not significantly different between undiagnosed males and females (OR = 1.03, p = 0.29, Fisher’s exact test). This suggests the existence of subtle sex differences in the genetic architecture of DD, especially with regard to known and undiscovered disorders. This could, for example, include sex-biased contribution of polygenic and/or environmental causes of DDs.

## Characteristics of the novel DD-associated genes and disorders

Based on semantic similarity^22^ between Human Phenotype Ontology terms, patients with DNMs in the same novel DD-associated gene were less phenotypically similar to each other, on average, than patients with DNMs in a consensus gene (p = 2.3 × 10^−11^, Wilcoxon rank-sum test; **Fig. 2a; Supplementary Figure 9**). This suggests that these novel disorders less often result in distinctive and consistent clinical presentations, which may have made these disorders harder to discover via a phenotype-driven analysis or recognise by clinical presentation alone. Each of these novel disorders requires a detailed genotype-phenotype characterisation, which is beyond the scope of this study.

**Figure 2:**
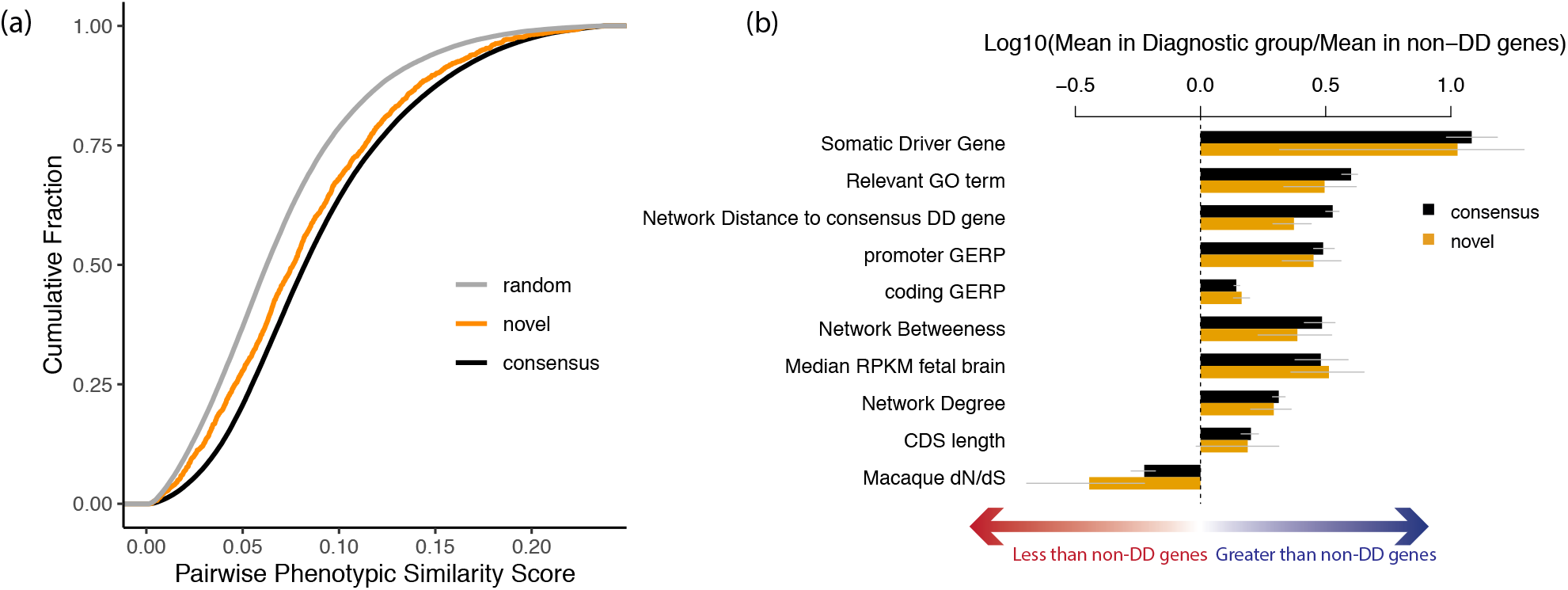
Functional properties and mechanisms of novel genes. (a) Comparing the phenotypic similarity of patients with DNMs in novel and consensus genes. Random phenotypic similarity was calculated from random pairs of patients. Patients with DNMs in the same novel DD-associated gene were less phenotypically similar than patients with DNMs in a known DD-associated gene (p = 2.3 × 10^−11^, Wilcoxon rank-sum test). (b) Comparison of functional properties of consensus and novel DD genes. Properties were chosen as those known to be differential between consensus and non-DD genes.

Overall, novel DD-associated genes encode proteins that have very similar functional and evolutionary properties to consensus genes, e.g. developmental expression patterns, network properties and biological functions (**Fig. 2b; Supplementary Table 5**). Despite the high-level functional similarity between known and novel DD-associated genes, the nonsynonymous DNMs in the more recently discovered DD-associated genes are much more likely to be missense DNMs, and less likely to be PTVs (discordant and novel; p = 1.2 × 10^−25^, chi-squared test). Fifteen of the 28 (54%) of the novel genes only had missense DNMs, and only a minority had more PTVs than missense DNMs. Consequently, we expect that a greater proportion of the novel genes will act via altered-function mechanisms (e.g. dominant negative or gain-of-function). For example, the novel gene *PSMC5* (DeNovoWEST p = 2.6 × 10^−15^) had one inframe deletion and nine missense DNMs, eight of which altered two structurally important amino acids that are both in the AAA+ ATPase domain within the 3D protein structure: p.Pro320Arg and p.Arg325Trp (**Supplementary Fig. 10a-b**), and so is likely to operate via an altered-function mechanism. None of the novel genes exhibited significant clustering of *de novo* PTVs.

We observed that missense DNMs were more likely to affect functional protein domains than other coding regions. We observed a 2.63-fold enrichment (p = 2.2 × 10^−68^, G-test) of missense DNMs residing in protein domains among consensus genes and a 1.80-fold enrichment (p = 8.0 × 10^−5^, G-test) in novel DD-associated genes, but no enrichment for synonymous DNMs (**Supplementary Table 6**). Four protein domain families in consensus genes were consistently enriched for missense DNMs (**Supplementary Table 7**): ion transport protein (PF00520, p = 6.9 × 10^−4^, G-test Bonferroni corrected), ligand-gated ion channel (PF00060, p = 4.0 × 10^−6^), protein kinase domain (PF00069, p = 0.043), and kinesin motor domain (PF00225, p = 0.027). Missense DNMs in all four enriched domain families have previously been associated with DD (**Supplementary Table 8**)^24–26^.

We observed a significant overlap between the 285 DNM-enriched DD-associated genes and a set of 369 previously described cancer driver genes^27^ (overlap of 70 genes; p = 1.7 × 10^−49^, logistic regression correcting for s_het_), as observed previously^28,29^, as well as a significant enrichment of nonsynonymous DNMs in these genes (**Supplementary Table 9**). This overlap extends to somatic driver mutations: we observe 117 DNMs at 76 recurrent somatic mutations observed in at least three patients in The Cancer Genome Atlas (TCGA)^30^. By modelling the germline mutation rate at these somatic driver mutations, we found that recurrent nonsynonymous mutations in TCGA are enriched 21-fold in the DDD cohort (p < 10^−50^, Poisson test, **Supplementary Fig. 11**), whereas recurrent synonymous mutations in TCGA are not significantly enriched (2.4-fold, p = 0.13, Poisson test). This suggests that this observation is driven by the pleiotropic effects of these mutations in development and tumourigenesis, rather than hypermutability.

## Recurrent mutations and potential new germline selection genes

We identified 773 recurrent DNMs (736 SNVs and 37 indels), ranging from 2-36 independent observations per DNM, which allowed us to interrogate systematically the factors driving recurrent germline mutation. We considered three potential contributory factors: (i) clinical ascertainment enriching for pathogenic mutations, (ii) greater mutability at specific sites, and (iii) positive selection conferring a proliferative advantage in the male germline, thus increasing the prevalence of sperm containing the mutation^31^. We observed strong evidence that all three factors contribute, but not necessarily mutually exclusively. Clinical ascertainment drives the observation that 65% of recurrent DNMs were in consensus genes, a 5.4-fold enrichment compared to DNMs only observed once (p < 10^−50^, proportion test). Hypermutability underpins the observation that 64% of recurrent *de novo* SNVs occurred at hypermutable CpG dinucleotides^32^, a 2.0-fold enrichment over DNMs only observed once (p = 3.3 × 10^−68^, chi-square test). We also observed a striking enrichment of recurrent mutations at the haploinsufficient DD-associated gene *MECP2*, in which we observed 11 recurrently mutated SNVs within a 500bp window, nine of which were G to A mutations at a CpG dinucleotide. *MECP2* exhibits a highly significant twofold excess of synonymous mutations within gnomAD^5^, suggesting that locus-specific hypermutability might explain this observation.

To assess the contribution of germline selection to recurrent DNMs, we initially focused on the 12 known germline selection genes, which all operate through activation of the RAS-MAPK signalling pathway^33,34^. We identified 39 recurrent DNMs in 11 of these genes, 38 of which are missense and all of which are known to be activating in the germline (see Supplement). As expected, given that hypermutability is not the driving factor for recurrent mutation in these germline selection genes, these 39 recurrent DNMs were depleted for CpGs relative to other recurrent mutations (6/39 vs 425/692, p = 3.4 × 10^−8^, chi-squared test).

Positive germline selection has been shown to be capable of increasing the apparent mutation rate more strongly^31^ than either clinical ascertainment (10-100X in our dataset) or hypermutability (~10X for CpGs). However, only a minority of the most highly recurrent mutations in our dataset are in genes that have been previously associated with germline selection. Nonetheless, several lines of evidence suggested that the majority of these most highly recurrent mutations are likely to confer a germline selective advantage. Based on the recurrent DNMs in known germline selection genes, DNMs under germline selection should be more likely to be activating missense mutations, and should be less enriched for CpG dinucleotides. **Table 1** shows the 16 *de novo* SNVs observed nine or more times in our DNM dataset, only two of which are in known germline selection genes (*MAP2K1* and *PTPN11*). All but two of these 16 *de novo* SNVs cause missense changes, all but two of these genes cause disease by an altered-function mechanism, and these DNMs were depleted for CpGs relative to all recurrent mutations. Two of the genes with highly recurrent *de novo* SNVs, *SHOC2* and *PPP1CB,* encode interacting proteins that are known to play a role in regulating the RAS-MAPK pathway, and pathogenic variants in these genes are associated with a Noonan-like syndrome^35^. Moreover, two of these recurrent DNMs are in the same gene *SMAD4*, which encodes a key component of the TGF-beta signalling pathway, potentially expanding the pathophysiology of germline selection beyond the RAS-MAPK pathway. Confirming germline selection of these mutations will require deep sequencing of testes and/or sperm^34^.

**Table 1:** Recurrent Mutations. *De novo* single nucleotide variants with more than 9 recurrences in our cohort annotated with relevant information, such as CpG status, whether the impacted gene is a known somatic driver or germline selection gene, and diagnostic gene group (e.g. consensus). “Recur” refers to the number of recurrences. “Likely mechanism” refers to mechanisms attributed to this gene in the published literature.

| Symbol | Chr | Position | Ref | Alt | Consequence | Recur | Likely mechanism | CpG | Somatic Driver Gene | Germline Selection Gene | DD status |
| --- | --- | --- | --- | --- | --- | --- | --- | --- | --- | --- | --- |
| PACS1 | 11 | 65978677 | C | T | missense | 36 | activating | Yes | - | - | consensus |
| PPP2R5D | 6 | 42975003 | G | A | missense | 22 | dominant negative | - | - | - | consensus |
| SMAD4 | 18 | 48604676 | A | G | missense | 21 | activating | - | Yes | - | consensus |
| PACS2 | 14 | 105834449 | G | A | missense | 13 | dominant negative | Yes | - | - | discordant |
| MAP2K1 | 15 | 66729181 | A | G | missense | 11 | activating | - | Yes | Yes | consensus |
| PPP1CB | 2 | 28999810 | C | G | missense | 11 | all missense/in frame | - | - | - | consensus |
| NAA10 | X | 153197863 | G | A | missense | 11 | all missense/in frame | Yes | - | - | consensus |
| MECP2 | X | 153296777 | G | A | stop gain | 11 | loss of function | Yes | - | - | consensus |
| CSNK2A1 | 20 | 472926 | T | C | missense | 10 | activating | - | - | - | consensus |
| CDK13 | 7 | 40085606 | A | G | missense | 10 | all missense/in frame | - | - | - | consensus |
| SHOC2 | 10 | 112724120 | A | G | missense | 9 | activating | - | - | - | consensus |
| PTPN11 | 12 | 112915523 | A | G | missense | 9 | activating | - | Yes | Yes | consensus |
| SMAD4 | 18 | 48604664 | C | T | missense | 9 | activating | Yes | Yes | - | consensus |
| SRCAP | 16 | 30748664 | C | T | stop gain | 9 | dominant negative | Yes | - | - | consensus |
| FOXP1 | 3 | 71021817 | C | T | missense | 9 | loss of function | Yes | - | - | consensus |
| CTBP1 | 4 | 1206816 | G | A | missense | 9 | dominant negative | Yes | - | - | discordant |

## Evidence for incomplete penetrance and pre/perinatal death

Nonsynonymous DNMs in consensus or significant DD-associated genes accounted for half of the exome-wide nonsynonymous DNM burden associated with DD (**Fig. 1b**). Despite our identification of 285 significantly DD-associated genes, there remains a substantial burden of both missense and protein-truncating DNMs in unassociated genes (those that are neither significant in our analysis nor on the consensus gene list). The remaining burden of protein-truncating DNMs is greatest in genes that are intolerant of PTVs in the general population (**Supplementary Fig. 12**) suggesting that more haploinsufficient (HI) disorders await discovery. We observed that PTV mutability (estimated from a null germline mutation model) was significantly lower in unassociated genes compared to DD-associated genes (p = 4.5 × 10^−68^, Wilcox rank-sum test **Fig. 3a**), which leads to reduced statistical power to detect DNM enrichment in unassociated genes. This is consistent with our hypothesis that many more HI disorders await discovery.

**Figure 3:**
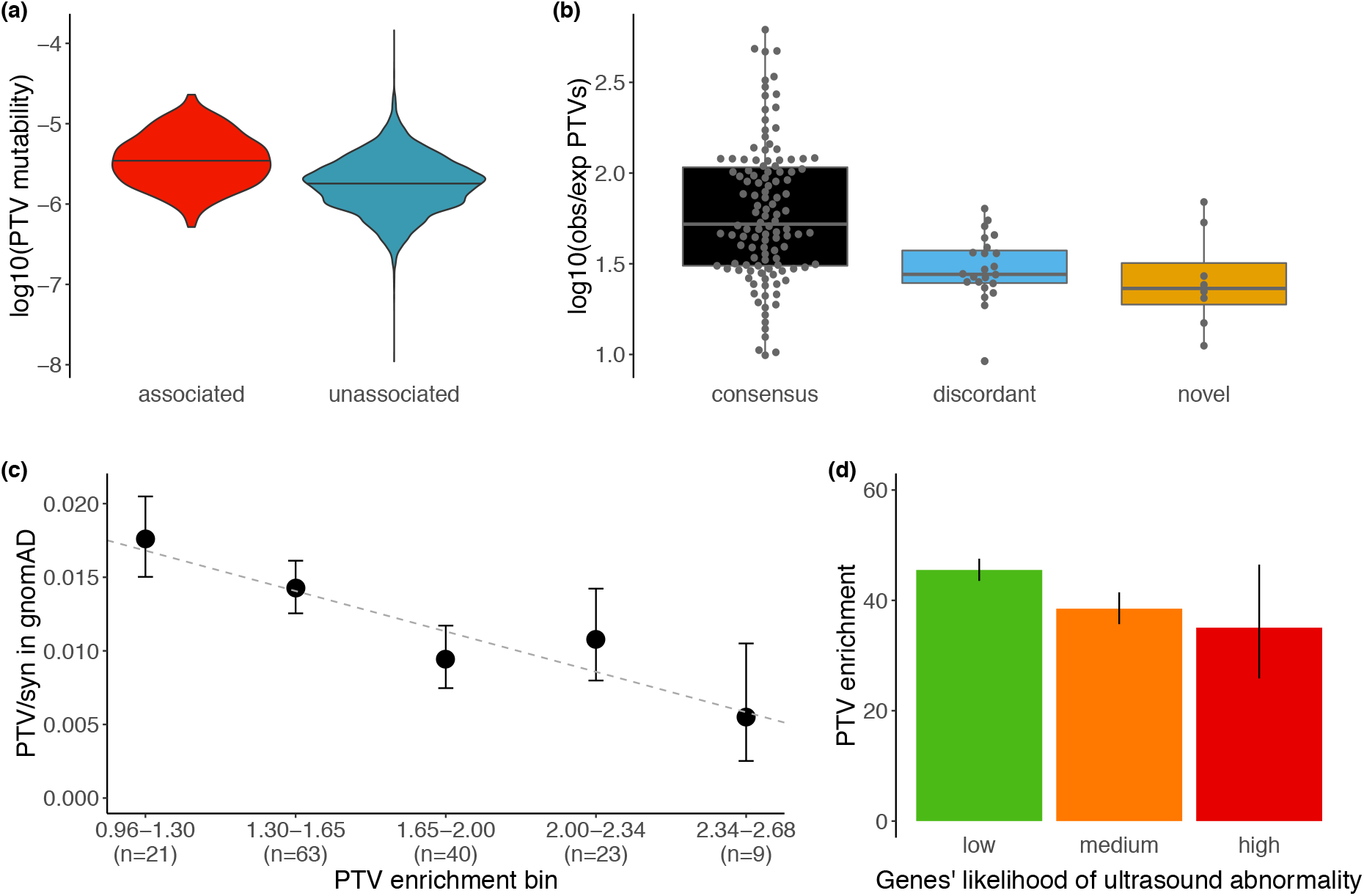
Impact of pre/perinatal death and penetrance on power. (a) PTV mutability is significantly lower in genes that are not significantly associated to DD in our analysis (“unassociated”, coloured blue) than in DD-associated genes (“associated”, coloured red; p = 4.6 × 10^−68^, Wilcox rank sum test). (b) Distribution of PTV enrichment in significant, likely haploinsufficient, genes by diagnostic group. (c) Comparison of the PTV enrichment in our cohort vs the PTV to synonymous ratio found in gnomAD, for genes that are significantly enriched for the number of PTV mutations in our cohort (without any variant weighting). PTV enrichment is shown as log10(enrichment). There is a significant negative relationship (p = 0.031, weighted regression). (d) Overall *de novo* PTV enrichment (observed / expected PTVs) across genes grouped by their clinician-assigned likelihood of presenting with a structural malformation on ultrasound during pregnancy. PTV enrichment is significantly lower for genes with a medium or high likelihood compared to genes with a low likelihood (p = 4.6 × 10^−5^, Poisson test).

A key parameter in estimating statistical power to detect novel HI disorders is the fold-enrichment of *de novo* PTVs expected in as yet undiscovered HI disorders. We observed that novel DD-associated HI genes had significantly lower PTV enrichment compared to the consensus HI genes (p = 0.005, Wilcox rank-sum test; **Fig. 3b**). Two additional factors that could lower DNM enrichment, and thus power to detect a novel DD-association, are reduced penetrance and increased pre/perinatal death, which here covers spontaneous fetal loss, termination of pregnancy for fetal anomaly, stillbirth, and early neonatal death. To evaluate incomplete penetrance, we investigated whether HI genes with a lower enrichment of protein-truncating DNMs in our cohort are associated with greater prevalences of PTVs in the general population. We observed a significant negative correlation (p = 0.031, weighted linear regression) between gene-specific PTV enrichment in our cohort and the gene-specific ratio of PTV to synonymous variants in gnomAD^5^, suggesting that incomplete penetrance does lower *de novo* PTV enrichment in individual genes in our cohort (**Fig. 3c**).

Additionally, we observed that the fold-enrichment of protein-truncating DNMs in consensus HI DD-associated genes in our cohort was significantly lower for genes with a medium or high likelihood of presenting with a prenatal structural malformation (p = 4.6 × 10^−5^, Poisson test, **Fig. 3d**), suggesting that pre/perinatal death decreases our power to detect some novel DD-associated disorders (see supplement for details).

## Modelling reveals hundreds of DD genes remain to be discovered

To understand the likely trajectory of future DD discovery efforts, we downsampled the current cohort and reran our enrichment analysis (**Fig. 4a**). We observed that the number of significant genes has not yet plateaued. Increasing sample sizes should result in the discovery of many novel DD-associated genes. To estimate how many haploinsufficient genes might await discovery, we modelled the likelihood of the observed distribution of protein-truncating DNMs among genes as a function of varying numbers of undiscovered HI DD genes and fold-enrichments of protein-truncating DNMs in those genes. We found that the remaining HI burden is most likely spread across ~1000 genes with ~10-fold PTV enrichment (**Fig. 4b**). This fold enrichment is three times lower than in known HI DD-associated genes, suggesting that incomplete penetrance and/or pre/perinatal death is much more prevalent among undiscovered HI genes. We modelled the missense DNM burden separately and also observed that the most likely architecture of undiscovered DD-associated genes is one that comprises over 1000 genes with a substantially lower fold-enrichment than in currently known DD-associated genes (**Supplemental Fig. 13**).

**Figure 4:**
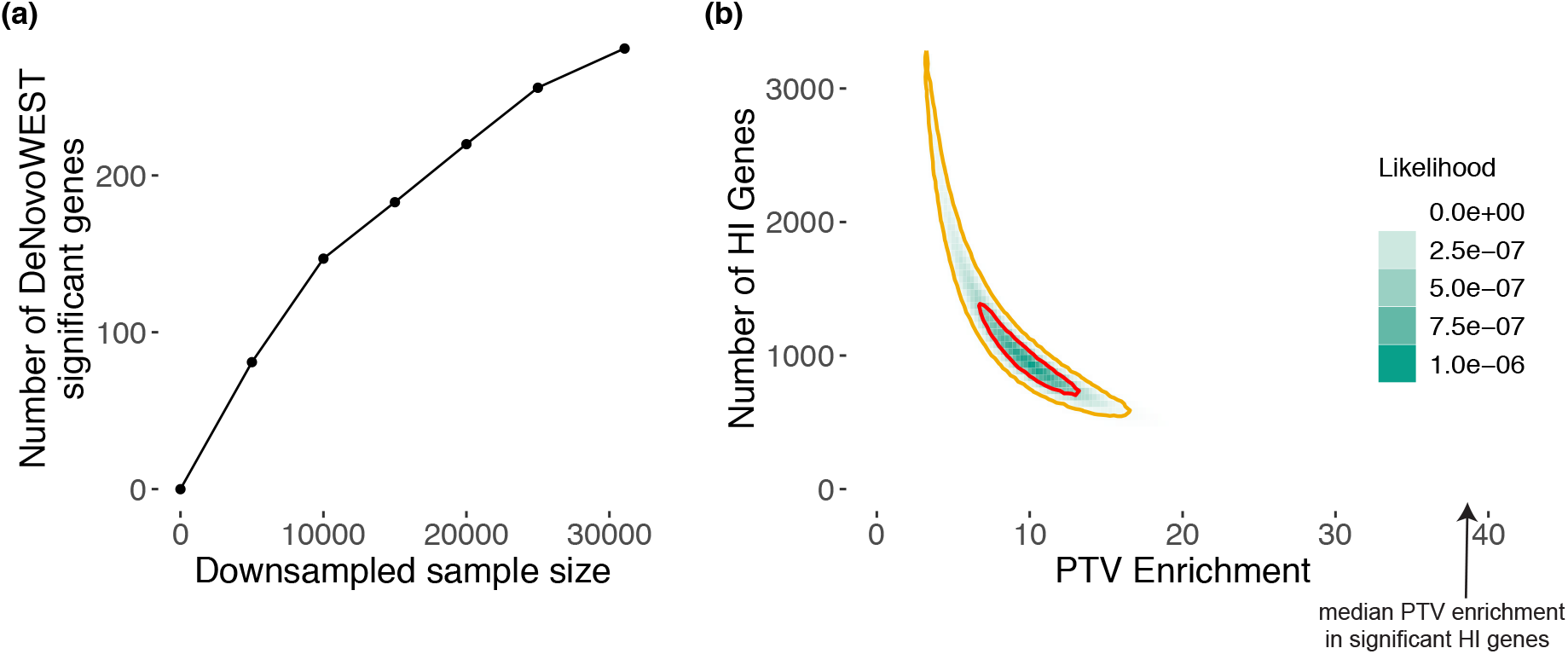
Exploring the remaining number of DD genes. (a) Number of significant genes from downsampling full cohort and running DeNovoWEST’s enrichment test. (b) Results from modelling the likelihood of the observed distribution of *de novo* PTV mutations. This model varies the numbers of remaining haploinsufficient (HI) DD genes and PTV enrichment in those remaining genes. The 50% credible interval is shown in red and the 90% credible interval is shown in orange. Note that the median PTV enrichment in significant HI genes (shown with an arrow) is 39.7.

We calculated that a sample size of ~350,000 parent-offspring trios would be needed to have 80% power to detect a 10-fold enrichment of protein-truncating DNMs for a gene with the median PTV mutation rate among currently unassociated genes. Using this inferred 10-fold enrichment among undiscovered HI genes, from our current data we can evaluate the likelihood that any gene in the genome is an undiscovered HI gene, by comparing the likelihood of the number of *de novo* PTVs observed in each gene to have arisen from the null mutation rate or from a 10-fold increased PTV rate. Among the ~19,000 non-DD-associated genes, ~1,200 were more than three times more likely to have arisen from a 10-fold increased PTV rate, whereas ~7,000 were three times more likely to have no *de novo* PTV enrichment.

## Discussion

In this study, we have discovered 28 novel developmental disorders by developing an improved statistical test for mutation enrichment and applying it to a dataset of exome sequences from 31,058 children with developmental disorders, and their parents. These 28 novel genes account for up to 1.0% of our cohort, and inclusion of these genes in diagnostic workflows will catalyse increased diagnosis of similar patients globally. We note that the value of this study for improving diagnostic yield extends well beyond these 28 novel genes; once newly validated discordant genes are included, the total number of genes added to the diagnostic workflows of the three participating centres ranged from 48-65 genes. We have shown that both incomplete penetrance and pre/perinatal death reduce our power to detect novel DDs postnatally, and that one or both of these factors are likely operating considerably more strongly among undiscovered DD-associated genes. In addition, we have identified a set of highly recurrent mutations that are strong candidates for novel germline selection mutations, which would be expected to result in a higher than expected disease incidence that increases dramatically with increased paternal age.

Our study represents the largest collection of DNMs for any disease area, and is approximately three times larger than a recent meta-analysis of DNMs from a collection of individuals with autism spectrum disorder, intellectual disability, and/or a developmental disorder^36^. Our analysis included DNMs from 24,348 previously unpublished trios, and we identified ~2.3 times as many significantly DD-associated genes as this previous study when using Bonferroni-corrected exome-wide significance (285 vs 124). In contrast to meta-analyses of published DNMs, the harmonised filtering of candidate DNMs across cohorts in this study should protect against results being confounded by substantial cohort-specific differences in the sensitivity and specificity of detecting DNMs.

Here we inferred indirectly that developmental disorders with higher rates of detectable prenatal structural abnormalities had greater pre/perinatal death. The potential size of this effect can be quantified from the recently published PAGE study of genetic diagnoses in a cohort of fetal structural abnormalities^37^. In this latter study, genetic diagnoses were not returned to participants during the pregnancy, and so the genetic diagnostic information itself could not influence pre/perinatal death. In the PAGE study data, 69% of fetal abnormalities with a genetically diagnosable cause died perinatally or neonatally, with termination of pregnancy, fetal demise and neonatal death all contributing. This emphasises the substantial impact that pre/perinatal death can have on reducing the ability to discover novel DDs from postnatal recruitment alone, and motivates the integration of genetic data from prenatal, neonatal and postnatal studies in future analyses.

To empower our mutation enrichment testing, we estimated positive predictive values (PPV) of each DNM being pathogenic on the basis of their predicted protein consequence, CADD score^3^, selective constraint against heterozygous PTVs across the gene (s_het_)^38^, and, for missense variants, presence in a region under selective missense constraint^4^. These PPVs should also be highly informative for variant prioritisation in the diagnosis of dominant developmental disorders. Further work is needed to see whether these PPVs might be informative for recessive developmental disorders, and in other types of dominant disorders. More generally, we hypothesise that empirically-estimated PPVs based on variant enrichment in large datasets will be similarly informative in many other disease areas.

We adopted a conservative statistical approach to identifying DD-associated genes. In two previous studies using the same significance threshold, we identified 26 novel DD-associated genes^1,39^. All 26 are now regarded as being diagnostic, and have entered routine clinical diagnostic practice. Had we used a significance threshold of FDR < 10% as used in Satterstrom, Kosmicki, Wang et al^40^, we would have identified 770 DD-associated genes. However, as the FDR of individual genes depends on the significance of other genes being tested, FDR thresholds are not appropriate for assessing the significance of individual genes, but rather for defining gene-sets. There are 184 consensus genes that did not cross our significance threshold in this study. It is likely that many of these cause disorders that were under-represented in our study due to the ease of clinical diagnosis on the basis of distinctive clinical features or targeted diagnostic testing. These ascertainment biases are, however, not likely to impact the representation of novel DDs in our cohort.

Our modelling also suggested that likely over 1,000 DD-associated genes remain to be discovered, and that reduced penetrance and pre/perinatal death will reduce our power to identify these genes through DNM enrichment. Identifying these genes will require both improved analytical methods and greater sample sizes. As sample sizes increase, accurate modelling of gene-specific mutation rates becomes more important. In our analyses of 31,058 trios, we observed evidence that mutation rate heterogeneity among genes can lead to over-estimating the statistical significance of mutation enrichment based on an exome-wide mutation model. We advocate the development of more granular mutation rate models, based on large-scale population variation resources, to ensure that larger studies are robust to mutation rate heterogeneity.

We anticipate that the variant-level weights used by DeNovoWEST will improve over time. As reference population samples, such as gnomAD^5^, increase in size, weights based on selective constraint metrics (e.g. s_het_, regional missense constraint) will improve. Weights could also incorporate more functional information, such as expression in disease-relevant tissues. For example, we observe that DD-associated genes are significantly more likely to be expressed in fetal brain (**Supplementary Fig. 14**). Furthermore, novel metrics based on gene co-regulation networks can predict whether genes function within a disease-relevant pathway^41^. As a cautionary note, including more functional information may increase power to detect some novel disorders while decreasing power for disorders with pathophysiology different from known disorders. Our analyses also suggest that variant-level weights could be further improved by incorporating other variant prioritisation metrics, such as upweighting variants predicted to impact splicing, variants in particular protein domains, or variants that are somatic driver mutations during tumorigenesis. In developing DeNovoWEST, we initially explored applying both variant-level weights and gene-level weights in separate stages of the analysis, however, subtle but pervasive correlations between gene-level metrics (e.g. s_het_) and variant-level metrics (e.g. regional missense constraint, CADD) presents statistical challenges to implementation. Finally, the discovery of less penetrant disorders can be empowered by analytical methodologies that integrate both DNMs and rare inherited variants, such as TADA^42^. Nonetheless, using current methods focused on DNMs alone, we estimated that ~350,000 parent-child trios would need to be analysed to have ~80% power to detect HI genes with a 10-fold PTV enrichment. Discovering non-HI disorders will need even larger sample sizes. Reaching this number of sequenced families will be impossible for an individual research study or clinical centre, therefore it is essential that genetic data generated as part of routine diagnostic practice is shared with the research community such that it can be aggregated to drive discovery of novel disorders and improve diagnostic practice.

## Acknowledgements

We thank the families and their clinicians for their participation and engagement. We are very grateful to our colleagues who assisted in the generation and processing of data. Inclusion of RadboudUMC data was in part supported by the Solve-RD project that has received funding from the European Union’s Horizon 2020 research and innovation programme under grant agreement No 779257. This work was in part financially supported by grants from the Netherlands Organization for Scientific Research: 917-17-353 to CG. The DDD study presents independent research commissioned by the Health Innovation Challenge Fund [grant number HICF-1009-003]. This study makes use of DECIPHER which is funded by Wellcome. See www.ddduk.org/access.html for full acknowledgement. The DDD study would like to acknowledge the tireless work of Rosemary Kelsell. Finally we acknowledge the contribution of an esteemed DDD clinical collaborator, M. Bitner-Glindicz, who died during the course of the study.

## Data Access

Sequence and variant level data and phenotypic data for the DDD study data are available through EGA study ID EGAS00001000775

RadboudUMC sequence and variant level data cannot be made available through EGA due to the nature of consent for clinical testing

GeneDx data cannot be made available through EGA due to the nature of consent for clinical testing. GeneDx has contributed deidentified data to this study to improve clinical interpretation of genomic data, in accordance with patient consent and in conformance with the ACMG position statement on genomic data sharing (see Supplementary Note for details).

Clinically interpreted variants and associated phenotypes from the DDD study are available through DECIPHER (https://decipher.sanger.ac.uk)

Clinically interpreted variants from RUMC are available from the Dutch national initiative for sharing variant classifications (https://www.vkgl.nl/nl/diagnostiek/vkgl-datashare-database) Clinically interpreted variants from GeneDx are deposited in ClinVar (https://www.ncbi.nlm.nih.gov/clinvar)

